# Persistent Post-Inflammatory Matrix Stiffening Defines a Premalignant Mechanical Niche in Ulcerative Colitis-Associated Neoplasia

**DOI:** 10.64898/2026.06.17.733043

**Authors:** Ziwei Wang, Ning Xie, Bing Xu, Jian Wu, Bo Cheng, Yutong Cheng, Haitao Shi, Qiuai Shu, Yingqi Li, Xiru Liang, Ameng Shi, Yuxin Peng, Bin Qin, Mingxuan Song, Kairuo Wang, Xin Liu, Jinhai Wang, Lu Li, Junye Liu, Na Liu, Feng Xu

**Affiliations:** Department of Gastroenterology, The Second Affiliated Hospital of Xi’an Jiaotong University, Xi’an 710004, P.R. China; The Key Laboratory of Biomedical Information Engineering of Ministry of Education, School of Life Science and Technology, Xi’an Jiaotong University, Xi’an 710049, P.R. China; Bioinspired Engineering and Biomechanics Center (BEBC), Xi’an Jiaotong University, Xi’an 710049, P.R. China; Department of Gastroenterology, Nanjing Drum Tower Hospital, Affiliated Hospital of Medical School, Nanjing University, Nanjing 210002, P.R. China; State Key Laboratory of Holistic Integrative Management of Gastrointestinal Cancers and National Clinical Research Center for Digestive Diseases, Xijing Hospital of Digestive Diseases, Fourth Military Medical University, Xi’an 710032, P.R. China; Department of Ultrasound Diagnosis, The Second Affiliated Hospital of Xi’an Jiaotong University, Xi’an 710004, P.R. China; Department of Pathology, The Second Affiliated Hospital of Xi’an Jiaotong University, Xi’an 710004, P.R. China; Hainan Medical University, Haikou 571199, P.R. China; Department of Radiation Medical Protection, School of Military Preventive Medicine, Air Force Medical University, Xi’an 710032, P.R. China; Department of Gastroenterology, Hainan General Hospital (Hainan Affiliated Hospital of Hainan Medical University), Haikou 570311, P.R. China

**Author notes:** These authors contributed equally to this work (Ziwei Wang, Ning Xie, Bing Xu, Jian Wu).

**Keywords:** ulcerative colitis, colitis-associated neoplasia, matrix stiffness, extracellular matrix remodeling, mechanotransduction, epithelial barrier dysfunction, premalignant epithelial remodeling

## Abstract

Patients with long-standing ulcerative colitis (UC) remain at risk for colorectal neoplasia even after overt inflammatory activity improves, suggesting that repaired mucosa may retain residual tissue-level abnormalities. Whether extracellular matrix remodeling leaves a persistent mechanical cue that contributes to early neoplastic remodeling is unknown. Here we identify post-inflammatory matrix stiffening as a premalignant mechanical niche in UC-associated neoplasia. In human colonic biopsies, collagen remodeling and atomic-force-microscopy-based mucosal stiffness increased from non-UC controls to UC and UC-associated dysplasia. A stiffness-associated transcriptional program was also enriched in histologically non-dysplastic mucosa from patients with UC-associated neoplasia. In mouse models, colonic stiffness increased along the colitis-to-tumorigenesis axis and remained elevated during apparent recovery, when epithelial permeability, junctional protein loss, crypt proliferation and nuclear β-catenin accumulation persisted despite reduced disease activity. Stiffness-controlled intestinal epithelial cultures showed that stiff substrates were sufficient to disrupt ZO-1 organization and enhance β-catenin redistribution, particularly under TNF-α stimulation. Pharmacological matrix normalization with β-aminopropionitrile partially restored barrier organization and attenuated early epithelial remodeling *in vivo*. Single-cell profiling identified a stiffness-associated MMP7-positive/YAP-active epithelial state that was enriched in collagen-rich regions and reduced after matrix softening.

Inhibition of YAP or MMP7 attenuated stiffness-associated junctional disruption and β-catenin redistribution. These findings suggest that inflammatory recovery and mechanical recovery can be uncoupled, and that persistent mucosal stiffening provides a tissue-level mechanism by which chronically injured UC mucosa may remain biologically vulnerable to premalignant epithelial remodeling.

## INTRODUCTION

Ulcerative colitis (UC) is increasingly managed through treat-to-target strategies that prioritize biomarker control, endoscopic healing and, when feasible, histological improvement ^1,2^. These goals have transformed clinical care, but they do not eliminate the long-term risk of colitis-associated neoplasia. Patients with extensive or long-standing UC continue to face increased colorectal cancer risk, and cumulative histological inflammation remains one of the most reproducible predictors of advanced neoplasia ^3–8^. This persistent risk suggests that the chronically injured mucosa may retain biological abnormalities that are not fully captured by inflammatory activity at a single clinical time point.

Recent high-impact studies have reframed inflammation-associated tumorigenesis as a problem of durable tissue memory and spatial niche remodeling ^9–13^. Colitis can leave long-lived epigenetic memory in colonic stem cells after histological recovery, thereby lowering the threshold for tumor growth after oncogenic challenge ^14^. In other epithelial organs, inflammatory memory can limit subsequent damage while also promoting tumorigenesis ^10^. Single-cell atlases of inflammatory gut disease have revealed injury-associated epithelial metaplasia and altered tissue architecture ^15^, and lineage-tracing work has shown that inflammation can enable non-stem epithelial lineages to act as alternative origins of intestinal tumors ^16^. Together with studies of oncogene-driven niche remodeling, wound-healing programs and early fibrotic or extracellular-matrix-defined tumor-permissive microenvironments ^11–13,17^, these findings indicate that cancer risk can be encoded in the post-injury tissue field, not only in ongoing inflammation or epithelial mutations.

Extracellular matrix (ECM) remodeling is a major, but still underused, layer of this post-injury field ^18–20^. Fibrosis and collagen reorganization are prominent features of chronic IBD, and recent work emphasizes that mechanical forces generated by matrix-rich tissues can initiate, propagate and potentially be targeted in fibrotic disease ^18,19,21^. In the intestine, matrix stiffness and mechanosensing pathways regulate epithelial fate, stem-cell maintenance, differentiation and barrier function ^22–24^. YAP/TAZ-associated mechanotransduction, PIEZO-dependent sensing, integrin signaling and fetal-like repair programs provide plausible routes by which remodeled ECM could shape epithelial states during regeneration and tumor initiation ^23,25–28^. However, whether mucosal stiffening persists after apparent inflammatory recovery in UC, and whether this mechanical abnormality contributes to premalignant epithelial remodeling, remains unresolved ^14^.

Here we investigated whether matrix stiffening persists after apparent inflammatory improvement and contributes to premalignant epithelial remodeling in UC-associated neoplasia. We integrate human colonic biopsies, public transcriptomic data, mouse models, stiffness-controlled epithelial cultures, matrix-normalization intervention and single-cell/spatial analyses. This framework addresses three linked questions: whether mucosal stiffening is detectable as a field-level abnormality during UC-associated neoplastic progression; whether this stiffness persists after apparent inflammatory recovery; and whether stiffened collagen-rich environments stabilize epithelial remodeling programs associated with barrier disruption and β-catenin activation. We identify a matrix metalloproteinase-7 (MMP7)-positive/YAP-active epithelial state positioned within collagen-rich stiffened mucosa and show that matrix normalization or YAP/MMP7 inhibition partially restores epithelial organization.

## MATERIALS AND METHODS

### Patient selection

Patients were recruited from the Department of Gastroenterology, The Second Affiliated Hospital of Xi’an Jiaotong University between November 2024 and December 2025. Colonic mucosal biopsies were obtained during colonoscopy. To reduce regional heterogeneity, biopsies used for the core stiffness and imaging analyses were collected from the sigmoid colon whenever anatomically feasible. Non-UC control samples were obtained from macroscopically and histologically normal mucosa of patients undergoing colonoscopy for colorectal polyps. UC diagnosis was established according to the *Chinese clinical practice guideline on the management of ulcerative colitis (2023, Xi’an edition)*^29^. Patients with colorectal cancer, primary sclerosing cholangitis or previous extensive intestinal resection were excluded. Based on histopathological evaluation, patients were further classified into UC and UC-dysplasia groups (Supplementary Table 1). The study was approved by the Institutional Review Board of The Second Affiliated Hospital of Xi’an Jiaotong University, and informed consent was obtained from all participants.

### Public dataset analysis

Public transcriptomic datasets were obtained from the Gene Expression Omnibus (GEO) database. GSE37283 was used to evaluate stiffness-associated transcriptional activity in normal controls, UC samples, and non-dysplastic mucosa from UC patients with dysplasia. A stiffness-associated score was calculated using single-sample gene set enrichment analysis based on a previously reported stiffness-related gene signature^30^. The full gene list used for this analysis is provided in Supplementary Table 2. Analyses were performed in R version 4.4.1.

### Animal Models

Female C57BL/6J mice aged 6-8 weeks were used. Chronic colitis and colitis-associated tumorigenesis were induced using dextran sodium sulfate (DSS) or azoxymethane (AOM)/DSS, respectively. For early recovery experiments, tissues were collected during active injury and apparent recovery according to the experimental timelines shown in the figures. To attenuate collagen crosslinking and matrix stiffening, β-aminopropionitrile (BAPN; 100 mg/kg/day) was administered by oral gavage. Disease activity index (DAI) and colonoscopy were used to monitor disease progression. All animal procedures were approved by the Institutional Animal Care and Use Committee of Xi’an Jiaotong University.

### Disease activity assessment and intestinal permeability

DAI was monitored throughout the experimental period based on body weight loss, stool consistency, and hematochezia. Intestinal permeability was evaluated using fluorescein isothiocyanate (FITC)-dextran gavage followed by plasma fluorescence quantification.

### Tissue stiffness and collagen characterization

Human tissue stiffness was measured using atomic force microscopy (AFM), while mouse colon tissues and polyacrylamide (PAA) hydrogels were characterized by rotational rheometry. Collagen deposition and architecture were assessed using Masson’s trichrome staining, Sirius Red staining, and second harmonic generation (SHG) imaging.

### Preparation of Stiffness-Controllable PAA Gels

Stiffness-controlled polyacrylamide hydrogels were prepared by adjusting acrylamide and bis-acrylamide concentrations to generate substrates with Young’s moduli of 0.6 kPa and 9.6 kPa, representing soft and stiff matrix conditions, respectively. Gel surfaces were activated with Sulfo-SANPAH and coated with rat tail collagen type I to support epithelial adhesion before organoid seeding.

### 2.5D intestinal organoid culture and stimulation

The 2.5D intestinal organoid culture was established according to previously described protocols^22,28^. Intestinal organoids were mechanically dissociated from mature 3D cultures and seeded onto collagen-coated PAA hydrogels to establish a 2.5D culture system. Following recovery, organoids were stimulated with recombinant TNF-α to model inflammatory conditions. Pharmacological inhibition experiments were performed using Verteporfin or an MMP7 inhibitor to evaluate the role of YAP and MMP7.

### Single-cell RNA sequencing and bioinformatic analysis

Colonic tissues from control, AOM/DSS, and BAPN-treated mice were dissociated into single-cell suspensions and processed using the 10X Genomics Chromium platform. Raw sequencing data were aligned and quantified using CellRanger. Downstream analyses were performed using Seurat. After quality control, normalization, dimensionality reduction, and clustering, epithelial cells were extracted and reclustered to identify remodeling-associated epithelial subpopulations. Differential gene expression, pathway enrichment, QuSAGE analysis, trajectory inference using Monocle, and developmental plasticity analysis using CytoTRACE were performed to characterize epithelial states associated with the stiffened microenvironment.

### Immunofluorescence staining and image analysis

Multiplex immunofluorescence staining was performed on paraffin-embedded tissues and 2.5D organoids. Images were acquired using confocal microscopy. Quantitative image analysis was conducted using ImageJ/Fiji to evaluate epithelial barrier integrity, YAP and β-catenin localization, proliferative activity, and collagen deposition.

### Statistical analysis

Data are presented as mean ± SD. Statistical analyses were performed using GraphPad Prism (version 9.0) and R (version 4.4.1). Statistical methods were selected according to data distribution and variance characteristics. Trend analysis among ordered clinical groups was performed using the Jonckheere–Terpstra test. Correlations were evaluated using Pearson correlation analysis. A two-sided *P* < 0.05 was considered statistically significant.

## RESULTS

### Mucosal matrix stiffness increases during UC-associated neoplastic progression

To determine whether matrix stiffening is coupled to UC-associated neoplastic progression, we combined direct biophysical measurements of human colonic biopsies with transcriptome-derived stiffness scoring and mouse models of colitis-associated tumorigenesis (**Fig. 1A**). This strategy allowed us to examine matrix mechanics at three levels: tissue stiffness in patient samples, stiffness-associated gene activity in non-dysplastic mucosa, and progressive ECM remodeling in experimental colitis-associated tumorigenesis.

**Figure 1.**
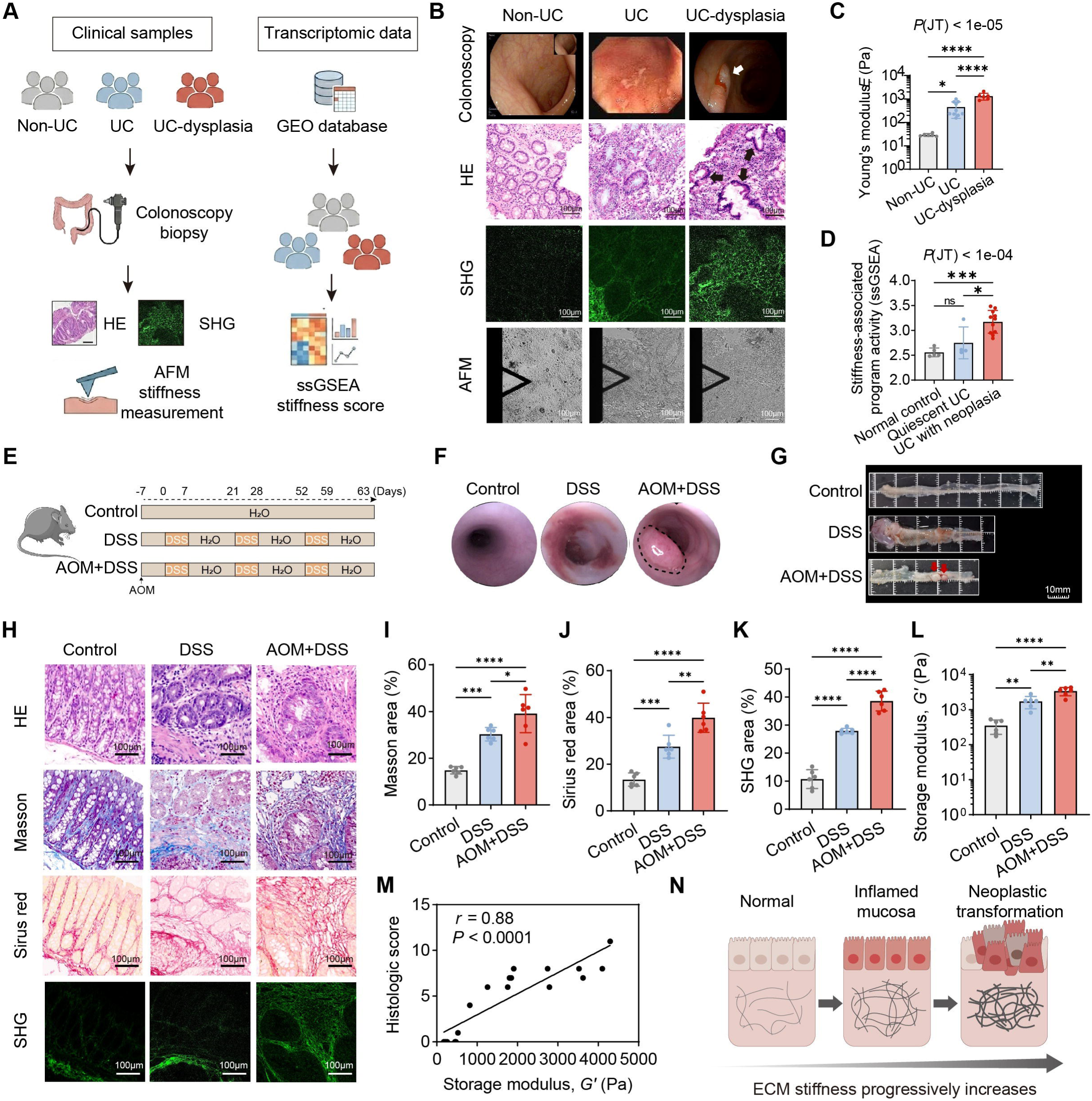
Matrix stiffness progressively increases during ulcerative colitis-associated tumorigenesis. (A) Workflow of colonic biopsy collection and stiffness characterization. Left, human biopsies analyzed by hematoxylin and eosin (H&E) staining, second harmonic generation (SHG) imaging, and atomic force microscopy (AFM). Right, transcriptomic stiffness scoring using a public GEO dataset. (B) Representative colonoscopic images, H&E staining, SHG imaging of fibrillar collagen, and AFM indentation images of human tissues from non-UC controls, UC, and UC-associated dysplasia. The white arrow indicates a dysplastic lesion under colonoscopy; black arrows indicate dysplastic colonic glands with nuclear stratification. Scale bars, 100 μm. (C) Tissue stiffness, measured as Young’s modulus (E), of human biopsies by AFM (Non-UC, *n* = 6; UC, *n* = 10; UC-associated dysplasia, *n* = 6). (D) Stiffness-associated transcriptomic scores calculated by single-sample gene set enrichment analysis (ssGSEA) using GSE37283 in normal controls, UC patients without neoplasia, and histologically non-dysplastic mucosa from UC patients with neoplasia. (E) Experimental design of DSS-induced colitis and azoxymethane/DSS (AOM/DSS)-induced colitis-associated cancer mouse models. (F, G) Representative colonoscopic (F) and gross morphological (G) images of mouse colons from control, DSS, and AOM/DSS groups. Scale bars, 10 mm. (H) Histological and extracellular matrix (ECM) characterization of mouse colons by H&E, Masson’s trichrome, Picrosirius Red, and SHG imaging. Scale bars, 100 μm. (I–K) Quantification of collagen deposition and ECM remodeling from (H), including Masson’s trichrome-positive area, Sirius Red-positive area, and SHG-positive area. (L) Quantification of tissue stiffness in mouse colons from control, DSS, and AOM/DSS groups, measured as storage modulus (G′) by rotational rheometry. (M) Pearson correlation analysis between tissue stiffness and histological score in mouse models. (N) Schematic summary illustrating progressive ECM remodeling and mechanical stiffening during UC-associated tumorigenesis. Data are shown as mean ± SD. Statistical methods are described in Methods. \**P* < 0.05, \*\**P* < 0.01, \*\*\**P* < 0.001, \*\*\*\**P* < 0.0001.

Human colonoscopic and histopathological analyses showed a continuum from non-UC control mucosa to inflamed UC mucosa and UC-associated dysplasia (**Fig. 1B**). SHG imaging revealed sparse pericryptal fibrillar collagen in non-UC mucosa, whereas collagen fibers progressively accumulated and thickened in UC and dysplastic tissues (**Fig. 1B; Supplementary Fig. 1A, B**). Consistent with these architectural changes, AFM showed a stepwise increase in mucosal Young’s modulus, with the highest stiffness in UC-associated dysplasia (**Fig. 1C**).

We next asked whether stiffness-associated abnormalities could be detected before overt dysplasia. In the GSE37283 dataset, histologically non-dysplastic mucosa from patients with UC-associated neoplasia showed higher stiffness-associated transcriptional activity than normal controls and quiescent UC samples without neoplasia (**Fig. 1D**). Because these samples were collected from macroscopically non-neoplastic regions, the result suggests that matrix-associated programs may mark a broader mucosal field rather than only the dysplastic lesion itself.

In mouse models, DSS-induced colitis and AOM/DSS-induced colitis-associated tumorigenesis produced progressive mucosal injury, collagen deposition and fibrillar remodeling (**Fig. 1E-H, Supplementary Fig. 1C, D**). Masson’s trichrome, Picrosirius Red and SHG quantification confirmed stepwise increases in collagen-rich ECM, and rotational rheometry demonstrated corresponding increases in bulk tissue storage modulus along the colitis-to-tumorigenesis axis (**Fig. 1I-L**). Tissue stiffness positively correlated with histological injury (*r* = 0.88, *P* < 0.0001; **Fig. 1M**). Together, these human and mouse data identify mucosal stiffening as a progressive and field-level feature of UC-associated neoplastic remodeling (**Fig. 1N**).

### Matrix stiffening persists during apparent inflammatory recovery and is associated with barrier dysfunction and early epithelial remodeling

We next asked whether matrix stiffening resolves when clinical inflammation improves. An early AOM/DSS model was used to capture the transition from active colitis to apparent post-inflammatory recovery (**Fig. 2A**). Disease activity increased during DSS exposure and declined toward baseline by day 21 (**Fig. 2B**). Despite this improvement, tissue stiffness remained significantly elevated at both day 7 and day 21 compared with controls (**Fig. 2C**).

**Figure 2.**
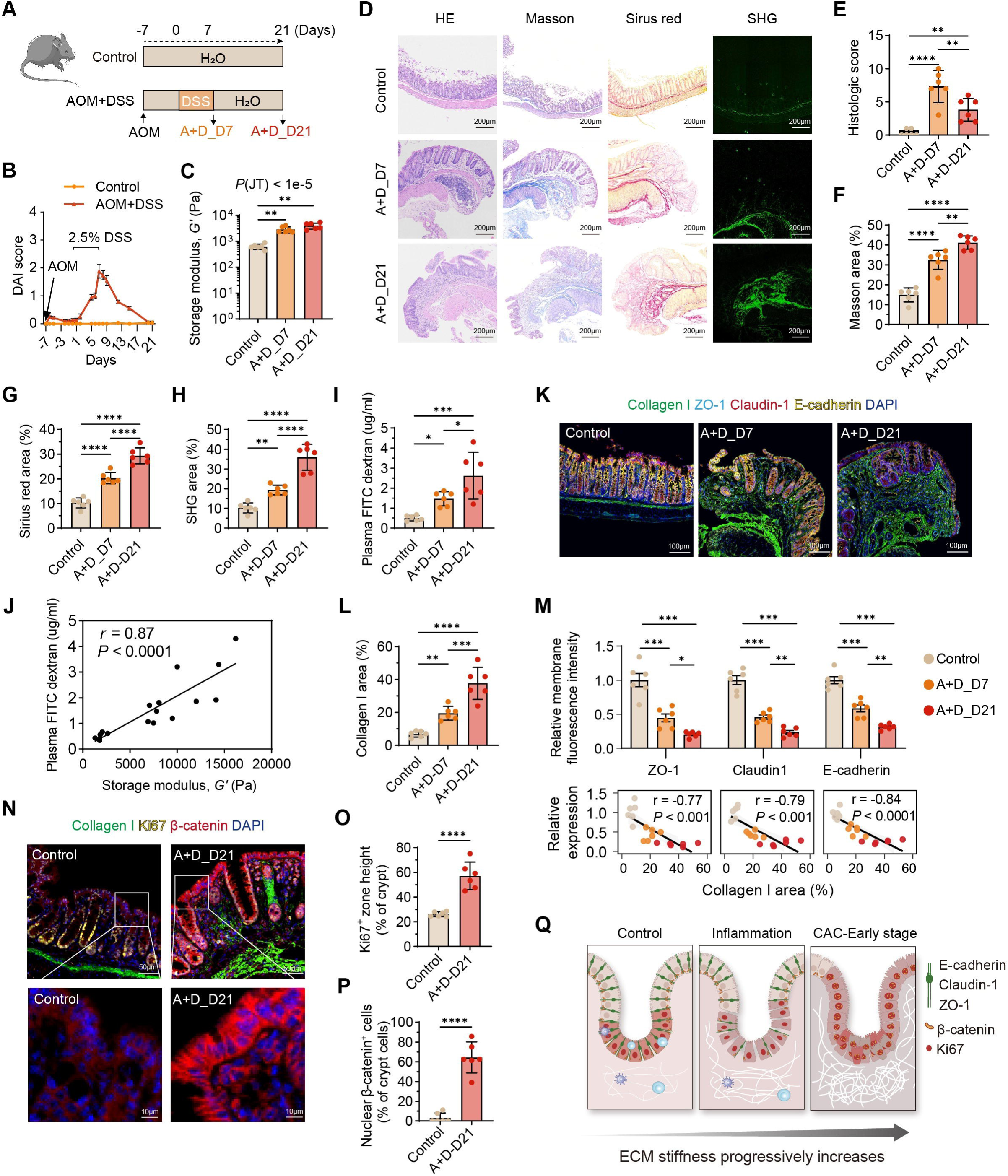
Persistent matrix stiffening during apparent post-inflammatory recovery is associated with epithelial barrier dysfunction and early epithelial remodeling. (A) Schematic of the early AOM/DSS model and tissue collection timeline. (B) Disease activity index (DAI) during AOM/DSS-induced colitis (*n* = 6). (C) Bulk colonic tissue stiffness, measured as storage modulus (G′), at day 7 (A+D_D7) and day 21 (A+D_D21) by rotational rheometry (*n* = 6). (D) Representative histological and ECM characterization of colonic tissues by H&E, Masson’s trichrome, Picrosirius Red, and SHG imaging. Scale bars, 200 μm. (E) Histological scores of control mice and AOM/DSS-treated mice at day 7 and day 21 (*n* = 6). (F–H) Quantification of ECM remodeling and collagen deposition based on Masson’s trichrome-positive area (F), Sirius Red-positive area (G), and SHG-positive area (H) (*n* = 6). (I) Plasma FITC-dextran levels indicating intestinal permeability (*n* = 6). (J) Pearson correlation analysis between tissue stiffness (G′) and intestinal permeability (plasma FITC-dextran). (K) Representative immunofluorescence images of Collagen I, tight junction proteins (ZO-1 and Claudin-1), adherens junction protein (E-cadherin), and DAPI in colonic sections. Scale bars, 100 μm. (L) Quantification of Collagen I-positive area. (M) Quantification of relative membrane fluorescence intensity of ZO-1, Claudin-1, and E-cadherin. Correlation plots show the relationship between Collagen I-positive area and junction protein expression. (N) Representative immunofluorescence staining for Collagen I, Ki67, β-catenin, and DAPI in control and AOM/DSS-treated mice at day 21. Scale bars, 50 μm (top) and 10 μm (bottom). (O, P) Quantification of Ki67-positive zone height (O) and percentage of epithelial cells with nuclear β-catenin localization (P). (Q) Schematic summary illustrating ECM stiffening-associated barrier disruption and early tumorigenic signaling. Data are shown as mean ± SD. Statistical methods are described in Methods. \**P* < 0.05, \*\**P* < 0.01, \*\*\**P* < 0.001, \*\*\*\**P* < 0.0001.

Histological injury was lower at day 21 than at day 7, but collagen deposition and fibrillar remodeling persisted. H&E, Masson’s trichrome, Picrosirius Red and SHG imaging showed residual mucosal remodeling during the apparent recovery phase (**Fig. 2D**). Quantification confirmed persistent increases in Masson’s trichrome-positive area, Sirius Red-positive area and SHG-positive collagen area at day 21 (**Fig. 2E-H**). Thus, inflammatory recovery and mechanical recovery were uncoupled in this model.

Persistent stiffening was accompanied by defective epithelial barrier organization. FITC-dextran permeability remained elevated during both active colitis and apparent recovery, and tissue stiffness positively correlated with plasma FITC-dextran levels (**Fig. 2I, J**). Multiplex immunofluorescence showed increased Collagen I together with reduced membrane localization of ZO-1, Claudin-1 and E-cadherin from day 7 through day 21 (**Fig. 2K-M**).

Epithelial remodeling also persisted after disease activity improved. At day 21, Ki67-positive epithelial cells expanded along the crypt axis and β-catenin showed increased nuclear localization (**Fig. 2N-P**). These data indicate that post-inflammatory matrix stiffening is not simply a passive scar, but a residual mucosal abnormality associated with barrier leakage, junctional loss and early premalignant epithelial signaling (**Fig. 2Q**).

### Stiff matrices are sufficient to disrupt epithelial junctions and promote premalignant remodeling in epithelial cultures

To test whether matrix stiffness can directly influence epithelial organization, we established stiffness-controlled 2.5D intestinal organoid cultures on collagen-coated polyacrylamide hydrogels (soft, 0.6 kPa; stiff, 9.6 kPa), with or without TNF-α stimulation (**Fig. 3A, Supplementary Fig. 1E**). This system isolates matrix stiffness from the cellular and biochemical complexity of inflamed tissue while preserving epithelial multicellular organization.

**Figure 3.**
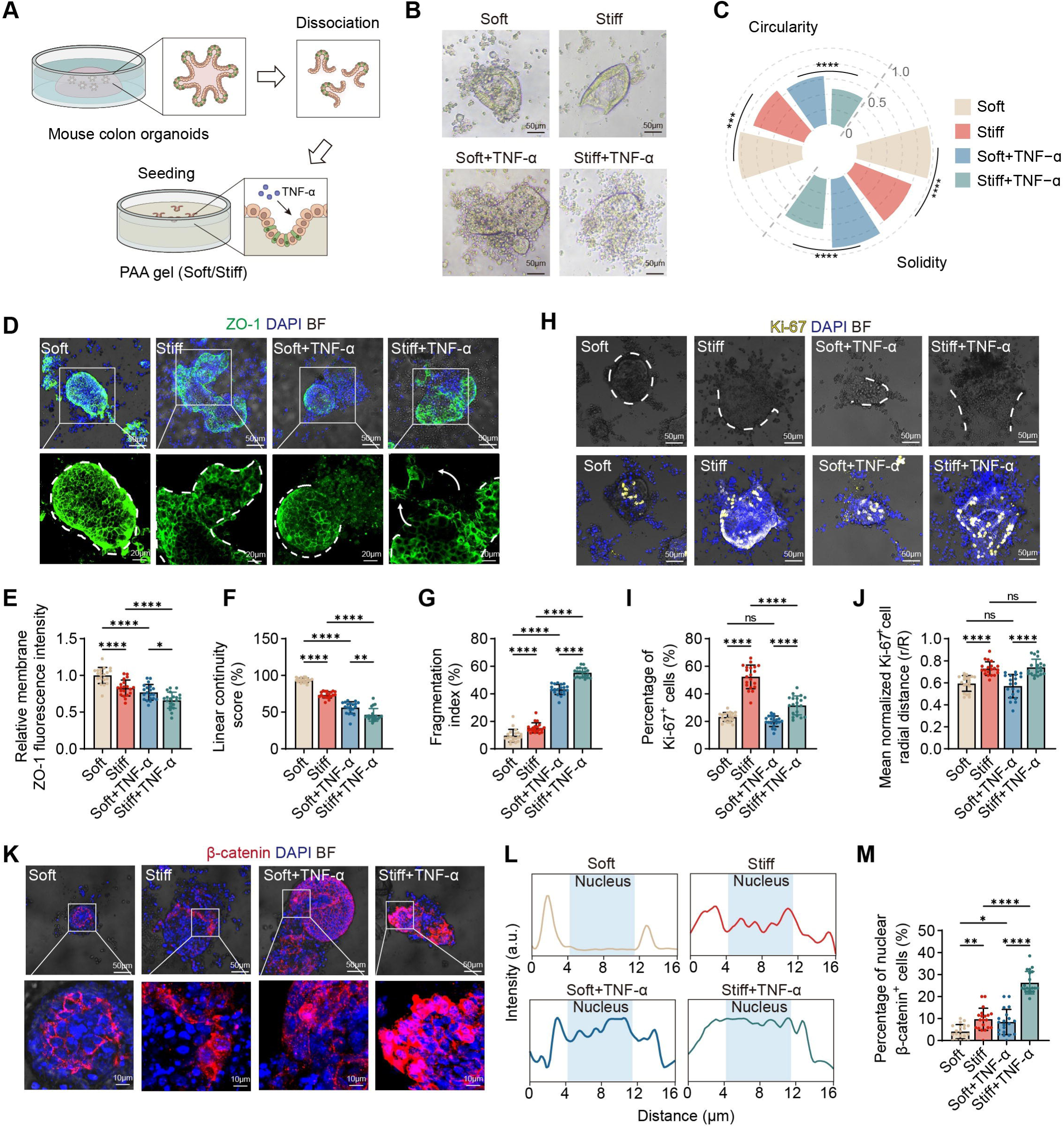
Matrix stiffness disrupts epithelial barrier organization and promotes dysplasia-associated remodeling in intestinal epithelial models. (A) Schematic illustration of the 2.5D intestinal organoid culture system. Primary mouse colonic organoids were dissociated and seeded onto stiffness-controlled polyacrylamide (PAA) hydrogels (soft, 0.6 kPa; stiff, 9.6 kPa), with or without TNF-α stimulation. (B, C) Representative bright-field images (B) and radar plot quantification (C) of organoid morphological parameters, including circularity and solidity. Scale bars, 50 μm. (D) Representative immunofluorescence staining of ZO-1 (green) overlaid with bright-field images in organoids cultured on soft or stiff matrices with or without TNF-α. Nuclei were counterstained with DAPI (blue). White dashed lines delineate organoid boundaries. Scale bars, 50 μm (top) and 20 μm (bottom). (E–G) Quantitative analysis of ZO-1 junctional integrity, including relative membrane fluorescence intensity (E), linear continuity score (F), and fragmentation index (G). (H) Representative immunofluorescence staining of Ki67 (yellow) overlaid with bright-field images in organoids cultured under the indicated conditions. Nuclei were counterstained with DAPI (blue). White dashed lines delineate organoid boundaries. Scale bars, 50 μm. (I, J) Quantification of Ki67-positive cells (I) and mean normalized radial distance (r/R) of Ki67-positive cells (J). (K) Representative immunofluorescence staining showing β-catenin (red) localization in organoids overlaid with bright-field images under the indicated conditions. Scale bars, 50 μm (top) and 10 μm (bottom). (L) Representative line-scan profiles of β-catenin fluorescence intensity across a single epithelial cell. The blue shaded area represents the nuclear region defined by DAPI. (M) Quantification of percentage of cells with nuclear β-catenin localization. *n* = 20 organoids per group from at least 3 independent experiments. Data are shown as mean ± SD. Statistical methods are described in Methods. ns, *P* > 0.05, \**P* < 0.05, \*\**P* < 0.01, \*\*\**P* < 0.001, \*\*\*\**P* < 0.0001.

Organoids cultured on stiff matrices showed altered morphology compared with those on soft matrices, and TNF-α amplified these effects (**Fig. 3B, C**). ZO-1 immunofluorescence revealed that stiff substrates reduced membrane enrichment and linear continuity of junctions while increasing junctional fragmentation, with the strongest disruption in stiff + TNF-α conditions (**Fig. 3D-G**).

Stiffness also promoted dysplasia-associated epithelial remodeling. Ki67-positive cells increased on stiff matrices and redistributed along the organoid radius, suggesting altered proliferative organization (**Fig. 3H-J**). β-catenin staining and line-scan analysis showed reduced membrane restriction and increased nuclear localization in stiff conditions, again accentuated by TNF-α (**Fig. 3K-M, Supplementary Fig. 1F**). These findings demonstrate that matrix stiffness is sufficient to perturb epithelial junctional architecture and cooperate with inflammatory stimulation to enhance premalignant epithelial features.

### Matrix normalization partially restores epithelial homeostasis in AOM/DSS-induced colitis

We next examined whether attenuating collagen crosslinking and matrix stiffening could improve epithelial organization *in vivo*. BAPN, an inhibitor of lysyl oxidase-mediated collagen crosslinking, was administered during the early AOM/DSS model (**Fig. 4A**). BAPN reduced colonic storage modulus compared with untreated AOM/DSS day 21 mice (**Fig. 4B**), supporting partial normalization of tissue mechanics.

**Figure 4.**
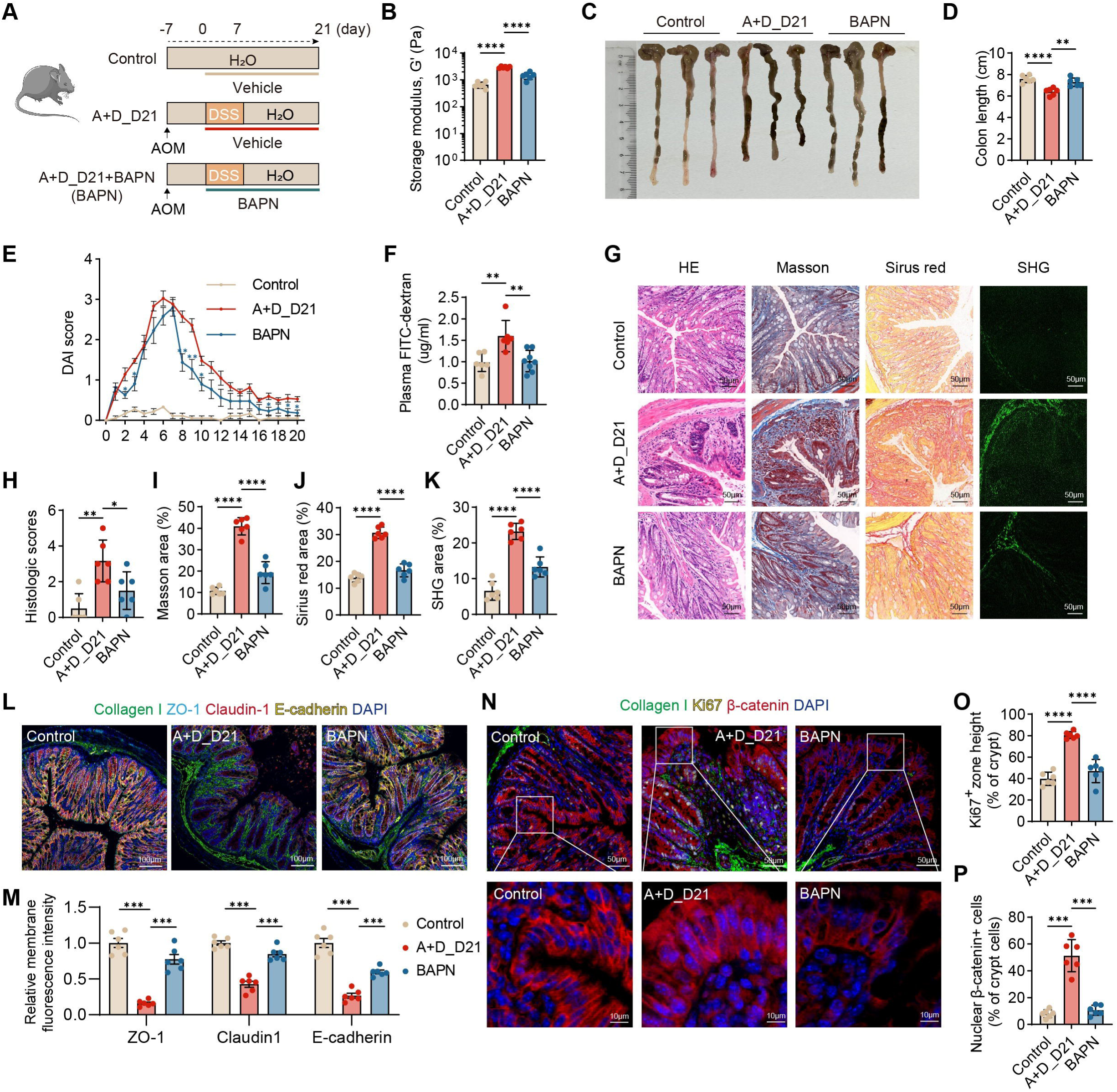
Matrix softening attenuates epithelial dysfunction and early dysplasia-associated remodeling in AOM/DSS-induced colitis. (A) Schematic of the experimental design for β-aminopropionitrile (BAPN) intervention in the AOM/DSS-induced colitis model. BAPN was administered to inhibit lysyl oxidase-mediated collagen crosslinking and subsequent matrix stiffening. (B) Quantification of colonic tissue stiffness, measured as storage modulus (G′), in control, AOM/DSS day 21 (A+D_D21), and BAPN-treated groups by rotational rheometry. (C, D) Representative gross images of colonic tissues (C) and quantification of colon length (D). (E) Disease activity index scores monitored during the experimental period. (F) Plasma FITC-dextran concentration measured at day 21 as an indicator of intestinal permeability. (G) Representative histological images of colonic sections stained with H&E, Masson’s trichrome, Picrosirius Red, and SHG imaging for collagen visualization. Scale bars, 50 μm. (H–K) Quantification of histological scores (H), Masson’s trichrome-positive area (I), Sirius Red-positive area (J), and SHG-positive area (K). (L, M) Representative immunofluorescence images (L) and quantification of relative membrane fluorescence intensity (M) of ZO-1, Claudin-1, and E-cadherin. Collagen I and DAPI are shown. Scale bars, 100 μm. (N) Representative immunofluorescence staining of Collagen I, Ki67, β-catenin, and DAPI. Scale bars, 50 μm (top) and 10 μm (bottom). (O, P) Quantification of Ki67-positive proliferative zone height (O) and percentage of epithelial cells with nuclear β-catenin localization (P). Data are shown as mean ± SD. Statistical methods are described in Methods. \**P* < 0.05, \*\**P* < 0.01, \*\*\**P* < 0.001, \*\*\*\**P* < 0.0001.

Matrix normalization improved several disease and barrier-associated readouts. BAPN-treated mice showed improved colon length, reduced disease activity during recovery, lower FITC-dextran permeability, reduced histological injury and decreased collagen-rich ECM deposition by Masson’s trichrome, Picrosirius Red and SHG analysis (**Fig. 4C-K**).

Epithelial junctional organization was also partially restored. ZO-1, Claudin-1 and E-cadherin membrane fluorescence increased after BAPN treatment (**Fig. 4L, M**). In parallel, Ki67-positive zone height and nuclear β-catenin-positive epithelial cells were reduced (**Fig. 4N-P**). Histopathological examination of major organs, including the heart, liver, spleen, lung, and kidney, revealed no overt structural abnormalities after chronic BAPN administration (**Supplementary Fig. 2**). Because BAPN acts at the tissue level, these data do not establish matrix stiffness as the only driver of epithelial remodeling. They do, however, support the functional relevance of collagen crosslinking and tissue mechanics in maintaining a post-inflammatory epithelial remodeling state.

### A rare MMP7-positive, YAP-active epithelial state emerges in collagen-rich stiffened mucosa

To define epithelial states associated with the stiffened mucosal microenvironment, we performed single-cell RNA sequencing of colonic tissues from control, AOM/DSS day 21 and BAPN-treated mice (**Fig. 5A**). Reclustering of epithelial cells identified major epithelial populations and a distinct MMP7-positive epithelial state enriched after AOM/DSS injury, and decreased after BAPN treatment (**Fig. 5B-D, Supplementary Fig. 3A-D**).

**Figure 5.**
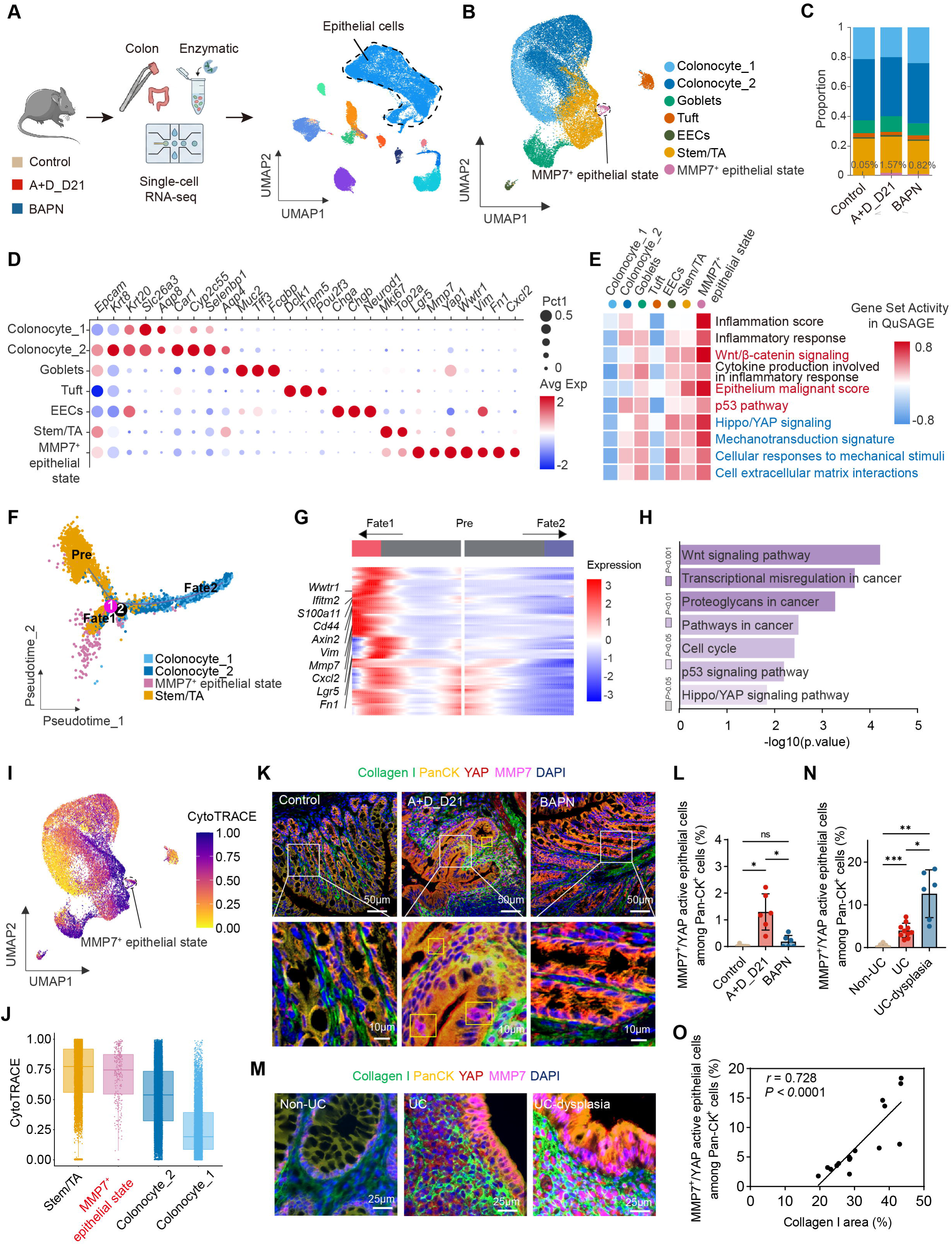
Single-cell transcriptomics identifies a stiffness-associated MMP7-positive/YAP-active epithelial state during colitis-associated tumorigenesis. (A) Experimental workflow for colonic tissue dissociation and single-cell RNA sequencing (scRNA-seq), with UMAP visualization showing the integrated landscape of all cell types from control, AOM/DSS day 21 (A+D_D21), and BAPN-treated groups. Epithelial cells are highlighted. (B) Reclustering of epithelial cells identified 7 epithelial subpopulations, including a distinct MMP7-positive/YAP-active epithelial population. (C) Relative proportions of epithelial subpopulations across groups. (D) Dot plot showing representative marker genes for epithelial subsets. The MMP7-positive/YAP-active epithelial population showed increased expression of *Mmp7*, *Yap1*, *Wwtr1*, *Vim*, *and Cxcl2* while retaining epithelial identity markers, including Epcam and Krt8. (E) Heatmap of QuSAGE gene set activity showing inflammatory, Wnt/β-catenin, malignant-associated, Hippo/YAP, mechanotransduction, cellular response to mechanical stimuli, and ECM-interaction programs across epithelial subsets. (F) Pseudotime trajectory analysis using Monocle, showing divergence from a progenitor state (Stem/TA, “Pre”) toward mature colonocytes or a MMP7-positive mechanosensitive epithelial branch. (G) Branched expression analysis modeling of epithelial cells. Heatmap showing dynamic expression of a gene module along the transition toward the MMP7-positive mechanosensitive epithelial branch. (H) KEGG pathway enrichment analysis of genes within this module. (I, J) CytoTRACE analysis projected on the UMAP (I) and quantified across epithelial subsets (J). (K) Representative immunofluorescence staining of pan-CK-positive/MMP7-positive/nuclear YAP-positive epithelial microdomains *in vivo*, showing localization near Collagen I-rich regions. These epithelial clusters were rare in control tissues, expanded in AOM/DSS-treated tissues, and reduced after BAPN-mediated matrix softening. Scale bars, 50 μm (top) and 10 μm (bottom). (L) Quantification of proportions of MMP7^+^ /YAP active epithelial cells among Pan-CK^+^ cells across groups. (M) Validation in human clinical specimens showing accumulation of pan-CK-positive/MMP7-positive/nuclear YAP-positive epithelial cells in UC and UC-associated dysplasia compared with non-UC controls. Scale bars, 25 μm. (N) Quantification of proportions of MMP7^+^ /YAP active epithelial cells among Pan-CK^+^ cells across groups. (O) Pearson correlation between collagen I-positive area (%) and MMP7□/YAP-active epithelial cells among Pan-CK□ cells (%) in human tissues from UC and UC-associated dysplasia. Data are shown as mean ± SD where applicable. Statistical methods are described in Methods. ns, *P* > 0.05, \**P* < 0.05, \*\**P* < 0.01, \*\*\**P* < 0.001, \*\*\*\**P* < 0.0001.

This epithelial state expressed *Mmp7* together with *Yap1*, *Wwtr1*, *Vim* and inflammatory chemokine-associated transcripts while retaining epithelial identity markers such as *Epcam* and *Krt8*. Gene set activity analysis showed enrichment of inflammatory response, Wnt/β-catenin, epithelial malignancy, Hippo/YAP, mechanotransduction and ECM-interaction programs (**Fig. 5E**). Pseudotime analysis placed this population on a remodeling-associated branch diverging from Stem/TA-like cells, and branched expression analysis identified modules enriched for Wnt signaling, cancer-associated pathways, cell cycle, p53 and Hippo/YAP signaling (**Fig. 5F-H**). CytoTRACE analysis indicated elevated developmental plasticity within the MMP7-positive epithelial state compared with mature colonocyte populations (**Fig. 5I, J**).

Spatial validation by multiplex immunofluorescence showed that MMP7-positive/nuclear YAP-positive epithelial microdomains localized near Collagen I-rich regions. These microdomains were rare in controls, expanded after AOM/DSS injury and were reduced after BAPN-mediated matrix normalization (**Fig. 5K, L**), and their abundance positively correlated with collagen I content (**Supplementary Fig. 3F**).

Human validation supported the relevance of this state to UC-associated neoplastic progression. Pan-CK-positive epithelial cells co-expressing MMP7 and nuclear YAP increased from non-UC controls to UC and UC-associated dysplasia (**Fig. 5M, N**), and similarly correlated with collagen I abundance in human tissues (**Fig. 5G**). Together, these data identify an MMP7-positive/YAP-active epithelial remodeling state positioned within collagen-rich stiffened mucosa.

### YAP and MMP7 inhibition attenuate stiffness-associated epithelial dysfunction

Finally, we tested whether YAP and MMP7 contribute functionally to stiffness-associated epithelial remodeling. In stiffness-controlled epithelial cultures, stiff substrates increased YAP fluorescence intensity and nuclear-to-cytoplasmic localization. YAP inhibition with Verteporfin reduced YAP activity and suppressed stiffness-induced MMP7 expression (**Fig. 6A-G**).

**Figure 6.**
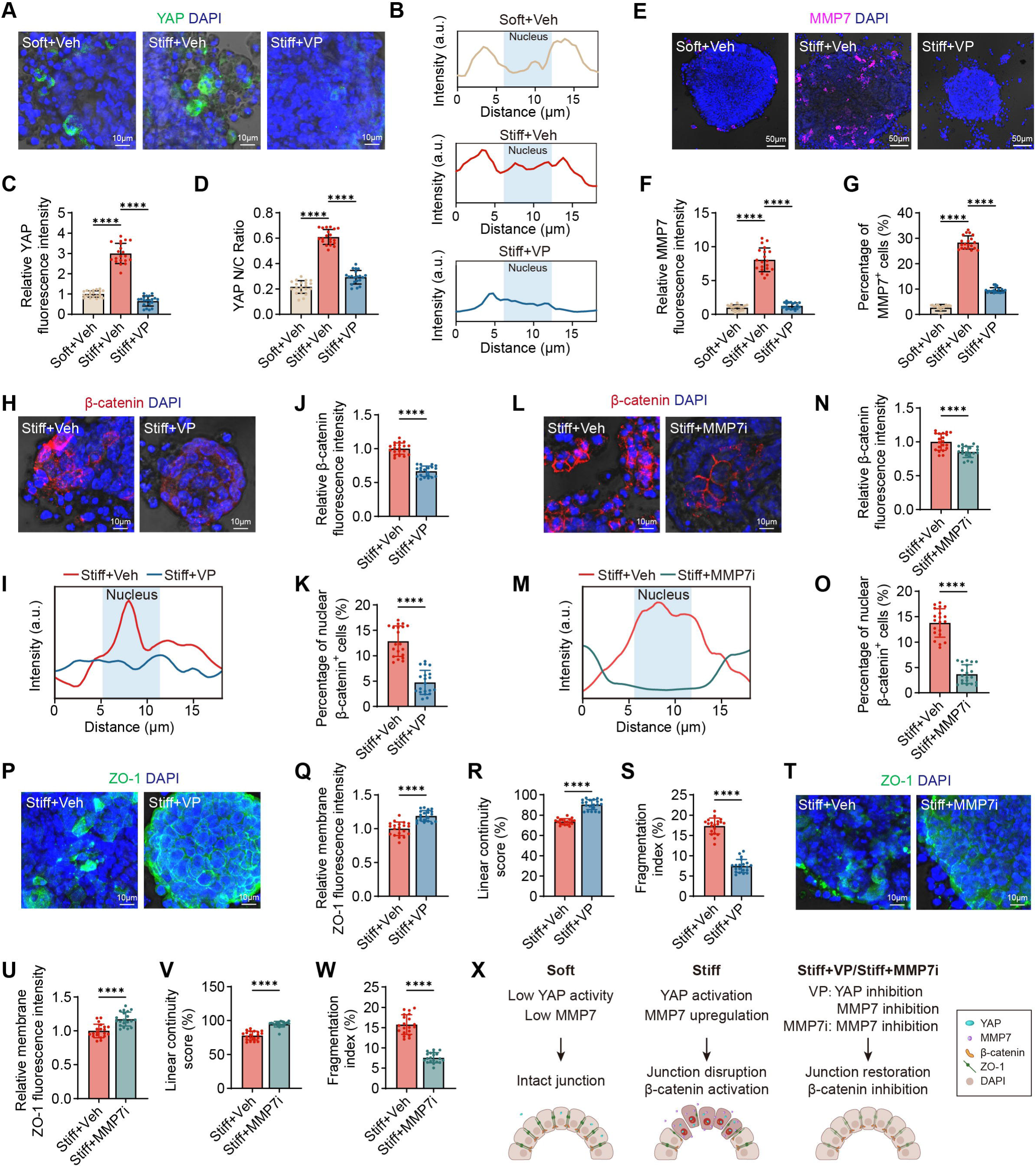
YAP and MMP7 inhibition attenuate stiffness-associated epithelial dysfunction in intestinal epithelial models. (A, B) Representative immunofluorescence images (A) and line-scan profiles (B) of YAP (green) in intestinal organoids cultured on soft, stiff, or stiff+Verteporfin (VP) substrates. Nuclei were counterstained with DAPI (blue). In (B), the shaded area indicates the DAPI-defined nucleus. Scale bars, 10 μm. (C, D) Quantification of relative YAP fluorescence intensity (C) and YAP nuclear-to-cytoplasmic ratio (D). (E) Representative immunofluorescence images of MMP7 (magenta) in organoids across soft, stiff, and stiff+VP groups. Scale bars, 50 μm. (F, G) Quantification of relative MMP7 fluorescence intensity (F) and percentage of MMP7-positive cells (G). (H, I) Representative immunofluorescence images (H) and line-scan profiles (I) of β-catenin (red) in stiff and stiff+VP groups. Scale bars, 10 μm. (J, K) Quantification of relative β-catenin fluorescence intensity (J) and percentage of cells with nuclear β-catenin localization (K) after YAP inhibition. (L, M) Representative immunofluorescence images (L) and line-scan profiles (M) of β-catenin (red) in stiff and stiff+MMP7 inhibitor (MMP7i) groups. Scale bars, 10 μm. (N, O) Quantification of percentage of cells with nuclear β-catenin localization (N) and relative β-catenin fluorescence intensity (O) after MMP7 inhibition. (P) Representative immunofluorescence images of ZO-1 (green) in stiff and stiff+VP groups. Scale bars, 10 μm. (Q–S) Quantification of ZO-1 organization after YAP inhibition, including relative membrane fluorescence intensity (Q), linear continuity score (R), and fragmentation index (S). (T) Representative immunofluorescence images of ZO-1 (green) in stiff and stiff+MMP7i groups. Scale bars, 10 μm. (U–W) Quantification of ZO-1 organization after MMP7 inhibition, including relative membrane fluorescence intensity (U), linear continuity score (V), and fragmentation index (W). (X) Schematic model illustrating that matrix stiffness promotes YAP activation and MMP7 upregulation, leading to junctional disruption and β-catenin nuclear accumulation, which are attenuated by pharmacologic inhibition of YAP or MMP7. *n* = 20 organoids per group from at least 3 independent experiments. Data are shown as mean ± SD. Statistical methods are described in Methods. \**P* < 0.05, \*\**P* < 0.01, \*\*\**P* < 0.001, \*\*\*\**P* < 0.0001.

YAP inhibition also attenuated premalignant epithelial remodeling. Verteporfin reduced β-catenin fluorescence intensity and nuclear β-catenin-positive cells in stiff cultures (**Fig. 6H-K**). It also improved ZO-1 membrane intensity, increased linear junctional continuity and reduced junctional fragmentation (**Fig. 6P-S**).

MMP7 inhibition produced a partially overlapping rescue phenotype. Blocking MMP7 reduced β-catenin redistribution and restored ZO-1 organization in stiff cultures (**Fig. 6L-O, T-W**). These perturbation experiments support a functional YAP/MMP7-associated module through which stiff matrix environments promote junctional disruption and β-catenin activation.

Consistent with these findings, similar protective effects of YAP or MMP7 inhibition were observed under TNF-α stimulation, as reflected by reduced β-catenin nuclear accumulation and improved ZO-1 junctional organization in organoids cultured on stiff matrices (**Supplementary Fig. 4**).

Rather than defining a strictly linear pathway, the data support a mechanically sensitive epithelial remodeling network in which YAP activity, MMP7 expression, inflammatory cues and β-catenin signaling reinforce barrier dysfunction and premalignant epithelial behavior (**Fig. 6X**).

## DISCUSSION

This study identifies persistent post-inflammatory matrix stiffening as a tissue-level abnormality that links chronic UC-associated injury to premalignant epithelial remodeling. Across human biopsies, public transcriptomic data, mouse models, stiffness-controlled epithelial cultures, matrix-normalization intervention and single-cell/spatial analyses, mucosal stiffening was coupled to collagen remodeling, epithelial junctional disruption, β-catenin redistribution and expansion of an MMP7-positive/YAP-active epithelial state. These findings support the concept that the repaired-appearing UC mucosa may remain mechanically abnormal even after overt inflammatory activity improves.

This study complements recent top-tier work on inflammatory and premalignant tissue memory. Colitis-induced epigenetic memory can persist in colonic stem cells long after tissue recovery and accelerate tumor growth after oncogenic mutation ^14^. Inflammatory injury can also drive epithelial metaplasia, noncanonical tumor origins and niche-dependent epithelial state transitions ^10,11,15,16^. Our data add a stromal-mechanical layer to this framework: repeated injury and repair can leave behind a stiffened collagen-rich mucosal field that provides aberrant mechanotransductive input even after inflammation declines. Thus, stiffness does not replace inflammation as a driver of colitis-associated tumorigenesis; rather, inflammation may generate a durable mechanical niche that continues to influence epithelial behavior.

A central implication is that inflammatory recovery and mechanical recovery can be dissociated. Current UC management appropriately emphasizes symptom control, biomarker normalization, endoscopic healing and histological improvement ^1,2,7^. However, clinical and molecular studies increasingly show that mucosal biology can remain perturbed during apparent remission ^14,31–34^. Our early AOM/DSS model extends this concept to tissue mechanics: disease activity and histological injury improved, yet collagen-rich ECM, stiffness, permeability defects and epithelial proliferative remodeling persisted. Partial reversal after BAPN treatment supports the view that residual stiffness is not merely a bystander scar, but a modifiable tissue feature that can continue to shape epithelial organization ^18–22,35^.

The single-cell and spatial data further refine this niche concept by linking stiffened mucosa to a defined epithelial state. The MMP7-positive/YAP-active population was not simply a diffuse injury marker: it retained epithelial identity, showed high plasticity, expressed inflammatory, Wnt/β-catenin, Hippo/YAP, mechanotransduction and ECM-interaction programs, localized near Collagen I-rich regions, expanded after AOM/DSS injury and contracted after matrix normalization. This is consistent with emerging models in which regeneration-associated or fetal-like epithelial programs can be adaptive during repair but maladaptive when stabilized in chronic injury settings ^15,16,25,36,37^. Positioning this state within collagen-rich mucosa helps convert a descriptive single-cell cluster into a spatially anchored premalignant mechanical niche.

Mechanistically, the data place YAP and MMP7 at a functional intersection between tissue mechanics and epithelial dysfunction. Matrix stiffness, collagen-mediated mechanotransduction, PIEZO-dependent sensing and YAP/TAZ signaling can regulate intestinal epithelial fate and injury-associated reprogramming ^22,23,26–28,35,38^. In our system, stiff substrates increased YAP nuclear localization and MMP7 expression, whereas YAP inhibition reduced MMP7 and improved epithelial junctional organization. MMP7 inhibition also attenuated β-catenin nuclear localization and restored ZO-1 continuity. Prior work supports links between stiff substrates, YAP-mediated MMP7 expression and MMP7-dependent barrier disruption ^39,40^. The most plausible model is not a single linear cascade, but a reinforcing network in which matrix stiffness, inflammatory cytokines, YAP activity, MMP7 and β-catenin signaling cooperate to destabilize epithelial homeostasis ^20,36,41^.

Clinically, these findings suggest that tissue mechanics could become an additional layer in future UC-associated neoplasia risk stratification. Existing risk assessment relies on disease duration, inflammatory burden, histology, endoscopic findings and emerging molecular markers . Our results suggest that collagen architecture, stiffness-associated transcriptional scores, mechanotransduction markers or imaging-derived ECM features may help identify mucosa that appears clinically improved but remains mechanically remodeled. Such measures should not be viewed as replacements for inflammatory control or dysplasia surveillance; rather, they may complement current approaches by capturing a physical component of residual mucosal risk.

Several limitations should be acknowledged. The human cohort is cross-sectional and modest in size, and longitudinal studies are required to determine whether mucosal stiffness independently predicts future dysplasia. The AOM/DSS model cannot fully separate prior inflammatory injury from matrix remodeling, although the persistence of stiffness after apparent recovery and partial reversal with BAPN support biological relevance. BAPN acts at the tissue level and does not define the responsible stromal cell population or mechanosensor. The 2.5D organoid system allowed controlled testing of stiffness but does not reproduce the full immune, stromal, microbial and biochemical complexity of the UC mucosa. Finally, although the YAP/MMP7 module was functionally supported, upstream mechanosensors and transcriptional partners remain to be resolved.

In conclusion, persistent post-inflammatory matrix stiffening defines a premalignant mechanical niche in UC-associated neoplasia. By sustaining barrier dysfunction and stabilizing an MMP7-positive/YAP-active epithelial state in collagen-remodeled mucosa, pathological stiffness provides one mechanism by which chronically injured tissue may remain biologically vulnerable after apparent inflammatory improvement. These findings argue that durable prevention of UC-associated neoplasia may require not only inflammatory control, but also restoration of the mechanical integrity of the mucosal microenvironment.

## Abbreviations

AFM: atomic force microscopy
AOM: azoxymethane
DSS: dextran sodium sulfate
BAPN: β-aminopropionitrile
DAI: disease activity index
ECM: extracellular matrix
FITC: fluorescein isothiocyanate
GEO: Gene Expression Omnibus
H&E: hematoxylin and eosin staining
MMP7: matrix metalloproteinase-7
PAA: polyacrylamide
SHG: second harmonic generation imaging
TAZ: transcriptional co-activator with PDZ-binding motif
TNF-α: tumor necrosis factor-α
UC: ulcerative colitis
YAP: yes-associated protein
ZO-1: zonula occludens-1

## Acknowledgements

The authors sincerely thank Qiaole Huang, Jingtong Zhang, and Lin Cao from Hainan Medical University for their participation and assistance in this study. The authors also acknowledge the imaging core facility of the Biomedical Experimental Center of Xi’an Jiaotong University for technical support and access to imaging equipment.

## Conflict of Interest

The authors declare no conflict of interest.

## Contributors

Ziwei Wang, Ning Xie, Bing Xu, and Jian Wu contributed equally to this work. Feng Xu, Na Liu, Ning Xie and Ziwei Wang conceived and designed the study. Ning Xie, Na Liu, and Kairuo Wang acquired funding for the project. Ziwei Wang, Ning Xie, Bing Xu and Jian Wu established experimental models and performed the major experiments. Ziwei Wang, Ning Xie and Bo Cheng analysed and interpreted the data. Ning Xie, Ziwei Wang and Xiru Liang prepared the figures and visualisations. Bing Xu, Jian Wu, Yutong Cheng, Haitao Shi, Qiuai Shu, Yingqi Li, Xiru Liang, Ameng Shi, Yuxin Peng, Bin Qin, Mingxuan Song, Kairuo Wang and Xin Liu participated in sample collection, experimental procedures and data acquisition. Ziwei Wang, Yutong Cheng, Bing Xu and Jian Wu curated the data. Ning Xie and Ziwei Wang drafted the manuscript. Junye Liu, Na Liu, Feng Xu and Jian Wu critically revised the manuscript for important intellectual content. Na Liu, Feng Xu, Lu Li, Junye Liu and Jinhai Wang supervised the study. All authors reviewed and approved the final version of the manuscript.

## Funding

This work was supported by the National Natural Science Foundation of China (No. 82303812, No. 82403890 and No. 82560549) and the Undergraduate Research and Innovation Training Program of Hainan Medical University (RZ2500002177).

## Ethics approval

This study was approved by the Institutional Review Board of The Second Affiliated Hospital of Xi’an Jiaotong University (2023513), and informed consent was obtained from all participants. All animal procedures were approved by the Institutional Animal Care and Use Committee of Xi’an Jiaotong University (XJTUAE2024-258).

## Data Availability Statement

The single-cell RNA sequencing dataset has been deposited in the National Genomics Data Center under accession number PRJCA066769.

## Notes

### Competing Interest Statement

The authors have declared no competing interest.

