## Supplementary Information for "Persistent Post-Inflammatory Matrix Stiffening Defines a Premalignant Mechanical Niche in Ulcerative Colitis-Associated Neoplasia"

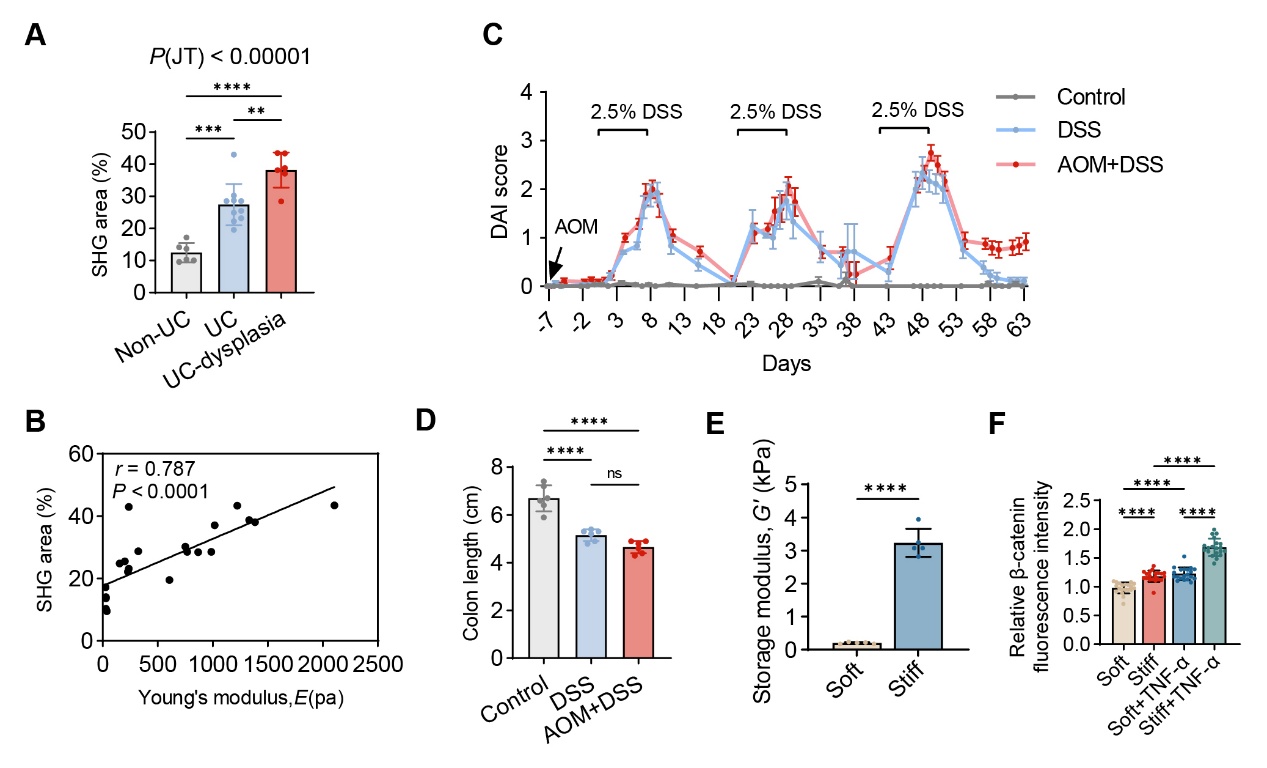

**Supplementary Figure 1. Additional data supporting Figure 1, Figure 2, and Figure 3.** (A) Quantification of second harmonic generation (SHG) area (%) in colonic tissues from non-UC controls, UC patients, and UC patients with dysplasia. (B) Pearson correlation analysis between SHG area (%) and tissue mechanical stiffness, represented by Young's modulus (Pa) (C) Disease Activity Index (DAI) scores monitored over the experimental period in Control, DSS, and AOM+DSS groups. (D) Colon length measurements at the experimental endpoint in Control, DSS, and AOM+DSS groups. (E) Storage modulus (G′) of soft and stiff polyacrylamide (PAA) gels measured by rotational rheometry. (F) Quantification of relative total β-catenin fluorescence intensity. Data are presented as mean ± SD. ***P* < 0.01, ****P* < 0.001, *****P* < 0.0001; *ns*, not significant.

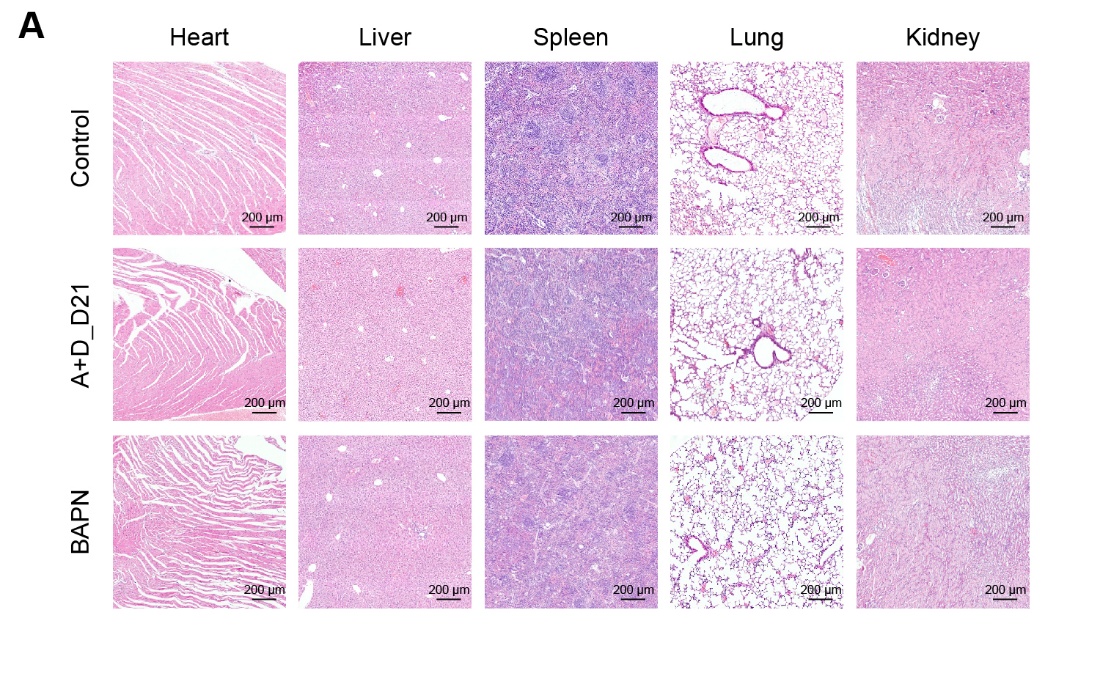

**Supplementary Figure 2. Systemic safety evaluation of BAPN intervention (Additional data supporting Figure 4).** Representative H&E-stained sections of the heart, liver, spleen, lung, and kidney from the Control, A+D_D21, and BAPN groups. No significant histopathological changes or systemic toxicities were observed across all major organs following BAPN administration. Scale bars, 200 μm.

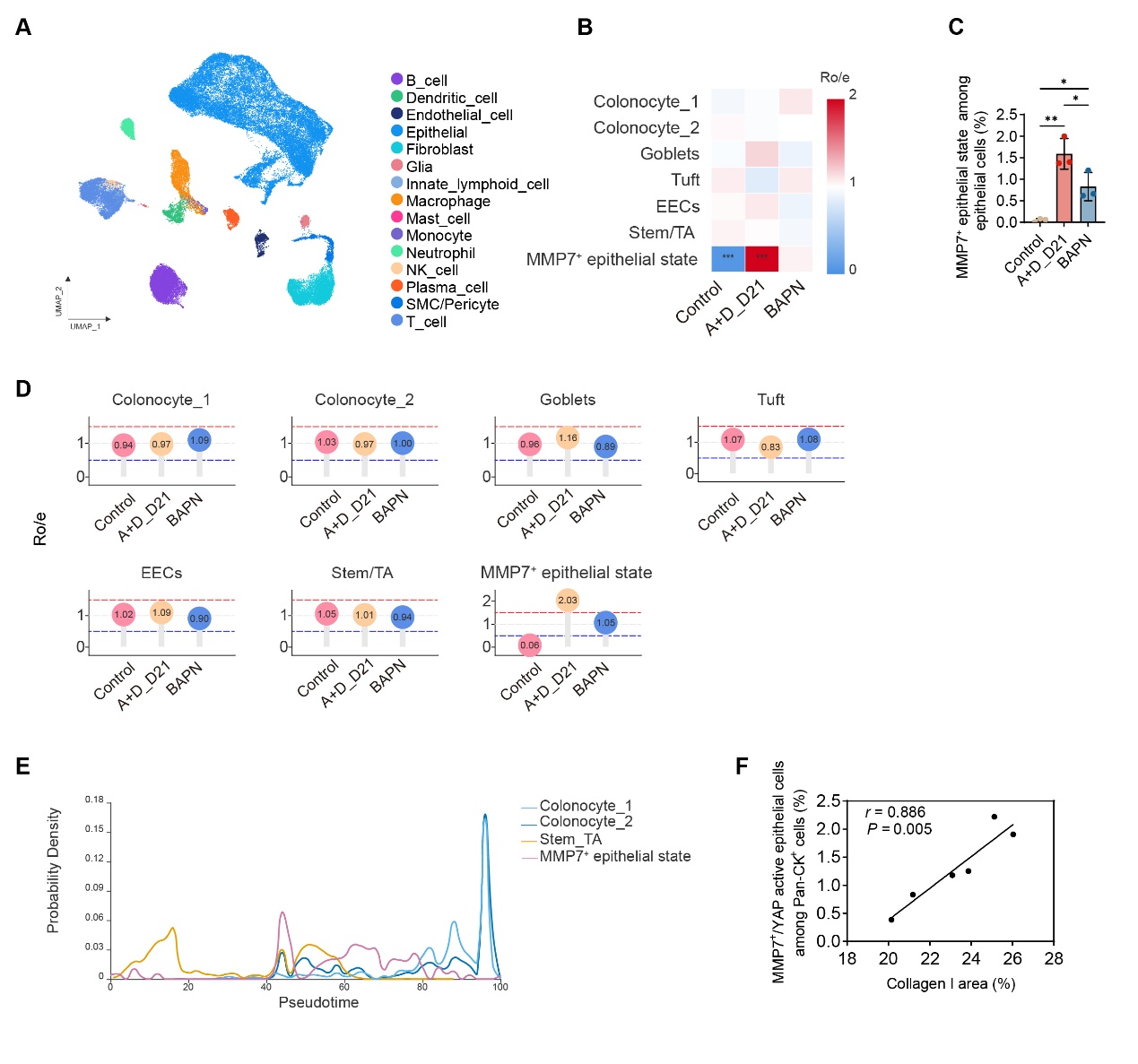

**Supplementary Figure 3. Additional analyses supporting the identification and dynamics of MMP7^+^ epithelial state during colitis-associated tumorigenesis (Additional data supporting Figure 5).** (A) UMAP visualization showing the integrated single-cell transcriptomic landscape of all major colonic cell types from the Control, AOM/DSS (A+D_D21), and BAPN groups. (B) Heatmap showing the ratio of observed to expected cell numbers (Ro/e) for seven epithelial subpopulations (Colonocyte_1, Colonocyte_2, Goblets, Tuft, EECs, Stem/TA, and MMP7^+^ epithelial state across the Control, A+D_D21, and BAPN groups. (C) Proportion of MMP7^+^ epithelial state among total epithelial cells in the Control, A+D_D21, and BAPN groups. (D) Ro/e analysis of individual epithelial subpopulations across experimental groups, demonstrating selective expansion of MMP7^+^ epithelial state in the A+D_D21 group and partial reversal following BAPN treatment. (E) Pseudotime probability density distributions generated by Monocle 3 trajectory analysis, illustrating the distribution of Colonocyte_1, Colonocyte_2, Stem/TA cells, and MMP7+ epithelial state populations along the epithelial differentiation trajectory. (F) Pearson correlation between collagen I-positive area (%) and MMP7⁺/YAP-active epithelial cells among Pan-CK⁺ cells (%) in A+D_D21 mice. **P* < 0.05, ***P* < 0.01

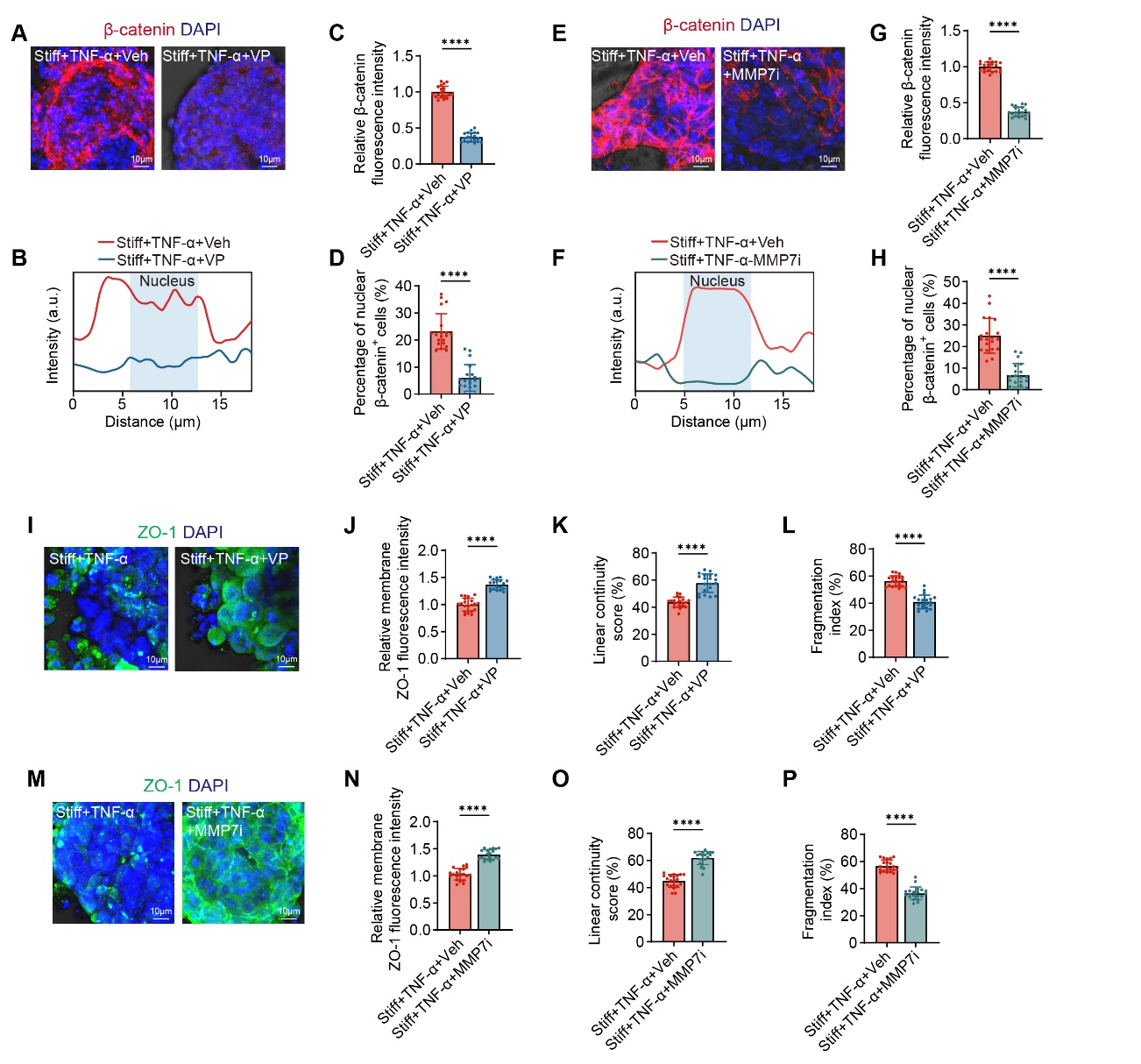

**Supplementary Figure 4. YAP and MMP7 inhibition attenuate stiffness-associated epithelial dysfunction in intestinal epithelial models with TNF-α (Additional data supporting Figure 6).** (A, B) Representative immunofluorescence images (A) and line-scan profiles (B) of β-catenin (red) in stiff+TNF-α+Veh and stiff+TNF-α+VP groups. Scale bars, 10 μm. (C, D) Quantification of relative β-catenin fluorescence intensity (C) and percentage of cells with nuclear β-catenin localization (D) after YAP inhibition. (E, F) Representative immunofluorescence images (E) and line-scan profiles (F) of β-catenin (red) in stiff+TNF-α+Veh and stiff+TNF-α+MMP7 inhibitor (MMP7i) groups. Scale bars, 10 μm. (G, H) Quantification of percentage of cells with nuclear β-catenin localization (G) and relative β-catenin fluorescence intensity (H) after MMP7 inhibition. (I) Representative immunofluorescence images of ZO-1 (green) in stiff+TNF-α+Veh and stiff+TNF-α+VP groups. Scale bars, 10 μm. (J–L) Quantification of ZO-1 organization after YAP inhibition, including relative membrane fluorescence intensity (J), linear continuity score (K), and fragmentation index (L). (M) Representative immunofluorescence images of ZO-1 (green) in stiff+TNF-α+Veh and stiff+TNF-α+MMP7i groups. Scale bars, 10 μm. (N–P) Quantification of ZO-1 organization after MMP7 inhibition, including relative membrane fluorescence intensity (N), linear continuity score (O), and fragmentation index (P). *n* = 20 organoids per group from at least 3 independent experiments. Data are shown as mean ± SD. *****P* < 0.0001.

**Supplementary Methods**

**Detailed DSS and AOM/DSS protocol**

For DSS-induced chronic colitis, mice received 2.5% (w/v) DSS (MP Biomedicals) in drinking water for three cycles, each consisting of 7 days DSS followed by 14 days regular water. In the AOM/DSS model, azoxymethane (AOM; Sigma-Aldrich) was administered intraperitoneally at 10 mg/kg seven days before DSS initiation. BAPN (MedChemExpress) was administered at 100 mg/kg/day by oral gavage. Control animals received vehicle (saline).

**Disease Activity Index (DAI) scoring**

The DAI was monitored daily to evaluate the severity of colitis during the AOM/DSS protocol. The score was calculated daily as the sum of scores for weight loss (0–4), stool consistency (0–4), and hematochezia (0–4). Specifically, weight loss was graded based on the percentage of initial body weight; stool consistency was evaluated by shape and adherence to the anus; and hematochezia was assessed by visual inspection, and with occult blood (score 1) detected using a fecal occult blood test kit (Leagene, TC0511). The average score (0–4) was used to indicate the severity of colitis.

**Intestinal Permeability Assay**

In vivo intestinal permeability was assessed using 4 kDa FITC-dextran (Beyotime, ST2930). Mice were fasted for 4 hours and then orally administered with FITC-dextran (600 mg/kg body weight). After 4 hours, peripheral blood was collected, and the plasma was separated by centrifugation at 3,000 rpm for 10 min. The concentration of FITC-dextran in the plasma was measured using a multi-mode microplate reader (SpectraMax M5) (excitation: 485 nm; emission: 535 nm).

**Atomic force microscopy (AFM) measurement**

Tissue preparation and AFM measurements were performed as previously described [1]. Biopsy or animal tissue samples were embedded in optimal cutting temperature (OCT) compound and cryosectioned. Serial sections were prepared, including a 4 μm section for hematoxylin and eosin (H&E) staining and histopathological evaluation, and a 20 μm section mounted on Superfrost Plus slides for AFM measurement. Sections were stored at −20 °C and measured within one week.

AFM measurements were performed using an NP-O10 cantilever (Bruker) equipped with a spherical polypropylene bead (radius: 6 μm; spring constant: ~0.06 N/m). For each sample, three independent regions were selected, and force–indentation curves were acquired over 8 × 8 maps within a 100 μm × 100 μm area in each region to account for tissue heterogeneity. The indentation speed was set at 2 μm/s with a maximum loading force of 2 nN.

The apparent Young’s modulus was calculated by fitting the force–indentation curves using the Hertz contact model, assuming a Poisson’s ratio of 0.5. The mean value from all measurements was used to represent the stiffness of each sample.

**Rotational Rheometry**

The storage modulus (*G'*) of mouse colon tissues and PAA gels was measured using MCR 302 rheometer (Anton Paar, Austria). Samples were placed between 8 mm parallel plates. Frequency sweep tests were performed at a constant strain of 1% (within the linear viscoelastic region) over a frequency range of 0.1–10 Hz at 37 °C. *G'* values at 1 Hz were used for comparison.

**2.5D Intestinal Organoid Culture and Stimulation**

The 2.5D intestinal organoid culture was established according to previously described protocols [2,3]. Briefly, mature 3D intestinal organoids were harvested from Matrigel and mechanically disaggregated by repeated pipetting in PBS containing calcium and magnesium (Servicebio). For each PAA gel, the quantity of organoid fragments derived from one Matrigel drop of a 24-well plate was seeded. After disaggregation, the organoid fragments were centrifuged at 100 × g for 3.5 minutes at room temperature, and the pellet was resuspended in organoid medium (Moji Bio). The organoids were then seeded in a small volume (50 μL) on top of the collagen-coated PAA gels and incubated at 37 °C for 2 hours to allow for initial attachment, after which an additional 550 μL of medium was added. Following 24–48 hours of recovery, organoids were exposed to recombinant mouse TNF-α (100 ng/mL; PeproTech) for an additional 24–48 hours to simulate an inflammatory microenvironment, and subsequently fixed for further immunofluorescence staining. To investigate the YAP-MMP7 axis, organoids on stiff substrates were treated with the YAP inhibitor Verteporfin (VP; 1 μM; MCE) or a selective MMP7 inhibitor (MMP7i; 10 μM; MCE) for 24-48 hours. Control groups received an equivalent concentration of DMSO (vehicle).

**Detailed Single-cell RNA sequencing and bioinformatic analysis**

Colonic tissues from the Control, AOM/DSS, and BAPN-treated groups were surgically removed and preserved in MACS Tissue Storage Solution (Miltenyi Biotec). The tissues were minced into small pieces (~1 mm^3^) on ice and enzymatically digested with collagenase I and DNase I for 45 minutes at 37 °C. The resulting cell suspension was sieved through a 70 μm strainer, treated with red blood cell lysis buffer , and further enriched using a MACS dead cell removal kit to ensure high cell viability (>85%). Single-cell scRNA-seq libraries were generated using the 10X Genomics Chromium Controller and Chromium Single Cell 3’ V3 Reagent Kits. Cells were loaded to generate Gel Bead-In-Emulsions (GEMs), followed by RT step, barcoded-cDNA amplification, and library construction. All libraries were sequenced on an Illumina Novaseq 6000 platform with 150 bp paired-end reads. Raw data were processed using CellRanger (v3.1.0) to align reads to the reference genome and generate feature-barcode matrices. Cells with over 200 expressed genes and a mitochondrial UMI rate below 10% were retained for further analysis. Seurat package was used for data normalization and UMAP-based clustering. Epithelial cells were extracted and re-clustered to identify distinct subpopulations. To delineate developmental transitions, single-cell trajectories were modeled using Monocle (utilizing DDR-Tree), and Branch Expression Analysis Modeling (BEAM) was applied to identify fate-determined gene modules at the trajectory bifurcation. Developmental plasticity was evaluated using the CytoTRACE algorithm to predict differentiation potential based on gene expression diversity. Additionally, the relative activation of mechanotransduction and pro-inflammatory pathways in epithelial clusters was quantified through QuSAGE (2.16.1) analysis. Functional implications of differentially expressed genes were further elucidated by Gene Ontology (GO) and KEGG pathway analyses.

**Detailed Immunofluorescence Staining**

Immunofluorescence staining was performed on both paraffin-embedded colonic tissue sections and 2.5D cultured organoids. For mouse and human colonic tissues, multiplex immunofluorescence staining was conducted using the multiplex staining kit (Huilan Biotech, RC0086plus-45RM) according to the manufacturer’s instructions. Briefly, sections were deparaffinized, rehydrated, and subjected to heat-induced antigen retrieval. To enable multi-color visualization, sequential rounds of primary antibody incubation, HRP-conjugated secondary antibody binding, and tyramide signal amplification (TSA) were performed. For 2.5D organoids, samples were fixed with 4% paraformaldehyde, permeabilized with 0.5% Triton X-100, and blocked with 5% BSA. Both tissues and organoids were incubated overnight at 4 °C with the following primary antibodies: Pan-CK (CST), MMP7 (CST), YAP (CST), Collagen I (CST), Ki67 (Servicebio), β-catenin (Abmart), E-cadherin (Proteintech), Claudin-1 (Proteintech) (all at 1:200 dilution), and ZO-1 (Proteintech, 1:500). Following primary incubation, samples were labeled with Alexa Fluor-conjugated secondary antibodies or TSA-fluorophores. Nuclei were counterstained with DAPI. Images were captured using an Olympus FV3000 confocal microscope. For 2.5D organoids, Z-stack images were captured and processed as maximum intensity projections. In representative panels, immunofluorescence channels were merged with bright-field (BF) images to provide a structural template for better delineation of organoid morphology.

**Image Quantification and Analysis**

Digital images were processed and quantified using ImageJ/Fiji software. For epithelial boundary and microdomain analysis, BF images were simultaneously acquired and used as structural templates to manually delineate Regions of Interest (ROIs). Subcellular localization of YAP and β-catenin was assessed using the Line Scanning tool; the nuclear-to-cytoplasmic (N/C) ratio was calculated by comparing the mean fluorescence intensity within the DAPI-defined nuclear area to that of the adjacent cytoplasmic region. Proliferative capacity was evaluated by measuring the Ki67^+^ zone height, defined as the distance from the crypt base to the uppermost Ki67^+^ cell normalized to total crypt height. For tight junction integrity, ZO-1 staining was quantified based on: (1) Relative Membrane Intensity; (2) Linear Continuity Score, representing the percentage of continuous ZO-1 signal along the junctional perimeter; and (3) Fragmentation Index, reflecting the frequency of junctional gaps or puncta per unit length. Collagen deposition was quantified by calculating the percentage of positive areas in Masson’s trichrome, Sirius Red, and SHG images. For clinical and animal samples, the percentage of nuclear β-catenin⁺ cells or MMP7⁺ cells were quantified in at least three randomly selected fields per sample, and the average value from these fields was used as a single biological replicate for statistical analysis.

**Detailed statistical analysis**

Normality was assessed using the Shapiro–Wilk test and homogeneity of variance using Levene’s test. Depending on data characteristics, Student’s t-test, Mann–Whitney U test, one-way ANOVA with Tukey’s test, Welch’s ANOVA with Games–Howell correction, or Kruskal–Wallis test with Dunn’s post hoc analysis were applied. Jonckheere–Terpstra analysis was used for ordered trend testing among Non-UC, UC, and UC-dysplasia groups. For animal experiments, *n* ≥ 6 mice per group were included, and quantitative measurements were averaged from at least three random fields or ten representative crypts per mouse.
